# Emerin regulation of nuclear stiffness is required for fast amoeboid migration in confined environments

**DOI:** 10.1101/2020.06.15.153007

**Authors:** Sandrine B. Lavenus, Karl W. Vosatka, Alexa P. Caruso, Maria F. Ullo, Ayesha Khan, Jeremy S. Logue

## Abstract

When metastasizing, tumor cells must traverse environments with diverse physicochemical properties. Recently, the cell nucleus has emerged as a major regulator of the transition from mesenchymal to fast amoeboid (leader bleb-based) migration. Here, in melanoma cells, we demonstrate that increasing nuclear stiffness through elevating Lamin A, inhibits fast amoeboid migration. Importantly, nuclei may respond to force through stiffening. A key factor in this process is the inner nuclear membrane (INM) protein, emerin. Accordingly, we determined the role of emerin in regulating fast amoeboid migration. Strikingly, we found that both the up- and down-regulation of emerin results in an inhibition of fast amoeboid migration. However, when key Src phosphorylation sites were removed, up-regulation of emerin no longer inhibited fast amoeboid migration. Interestingly, in confined cells, Src activity was low, as measured by a Src biosensor. Thus, the fast amoeboid migration of melanoma cells depends on the precise calibration of emerin activity.

**Summary Statement:** In mechanically constrictive microenvironments, amoeboid migrating melanoma cells require emerin for the precise calibration of nuclear stiffness.

## Introduction

Cell migration is integral to embryonic development, immune surveillance, and wound healing. When de-regulated, cell migration may also contribute to disease. In transformed cells, cell migration is an essential aspect of metastasis, which accounts for the vast majority of cancer deaths. Because the development of therapeutics that prevent or abate cancer metastasis depends on it, elucidating the molecular mechanisms that regulate the migration of cancer cells is an important problem.

It has become clear that cancer cells may utilize a number of mechanisms to disseminate from a primary tumor to distant tissues. This includes mesenchymal, collective, lobopodial, osmotic engine, and fast amoeboid (leader bleb-based) migration (Paul et al., 2017; Yamada and Sixt, 2019). Osmotic engine and leader bleb-based migration (LBBM) occurs specifically within confined environments, such as along or within micro-capillaries/lymphatics and between epithelial surfaces (Logue et al., 2015; Stroka et al., 2014). During LBBM, cells protrude a large, intracellular pressure driven, plasma membrane (PM) bleb (Liu et al., 2015; Logue et al., 2015; Ruprecht et al., 2015). Because cells migrate in the direction of this bleb, it was termed a ‘leader bleb’ (Logue et al., 2015). With non-specific friction, a rapid cortical actin flow in leader blebs provides the motive force for cell movement (Bergert et al., 2015). Because it only requires non-specific friction, cells may use LBBM in diverse physiochemical environments (Madsen and Sahai, 2010). This property is likely to promote distant metastasis. A phenotypically similar mode of migration has been observed in breast and melanoma tumors by intravital imaging (Tozluoglu et al., 2013). Notably, amoeboid migrating melanoma cells have also been observed at the invasive fronts of patient tumors (Cantelli et al., 2015).

Because cancer cells may adopt multiple mechanisms of migration, preventing or abating metastasis using therapeutics is challenging. For instance, we previously demonstrated that fast amoeboid migrating cells are resistant to Src inhibitors (Logue et al., 2018; Ullo and Logue, 2018). This is in contrast to mesenchymal migration, which requires Src activity to form focal adhesions (Webb et al., 2004). Therefore, the identification of factors required by all modes of cancer cell migration is necessary for the development of effective therapeutics. Within tissues, motile cells must deform in order to squeeze through confining environments. Therefore, factors that regulate cell deformability may be ideal therapeutic targets.

In recent years, it has become clear that the nucleus plays an integral role in sensing the cellular environment. In a confined cell, it has been demonstrated that the nuclear membrane becomes tensed, which together with increased intracellular calcium, recruits cytosolic Phospholipase A2 (cPLA2) (Lomakin et al., 2020; Venturini et al., 2020). At the nuclear membrane, cPLA2 liberates arachidonic acid to increase actomyosin contractility (Gong et al., 1992). Intracellular calcium is released upon rupture of the ER, which becomes compressed between the nucleus and PM in confined cells (Lomakin et al., 2020; Venturini et al., 2020). Thus, cPLA2 regulates the phenotypic transition from mesenchymal to fast amoeboid migration in confined environments.

Because of the essential role of nuclear membrane tension, factors that regulate the deformability of the nucleus are likely to be potent regulators of fast amoeboid migration. Accordingly, we set out to determine the role of the inner nuclear membrane (INM) protein, emerin, in regulating fast amoeboid migration. Emerin is a LEM (LAP2, emerin, MAN1) domain containing protein, which connects emerin to chromatin through a direct interaction with Barrier to Autointegration Factor (BAF) (Berk et al., 2013). Interestingly, it was demonstrated that emerin confers nuclear adaptation to force. More specifically, using magnetic beads coated with anti-Nesprin-1 antibody and isolated nuclei, repeated pulling by magnetic tweezers led to nuclear stiffening in an emerin dependent manner (Guilluy et al., 2014). Additionally, this effect was found to depend on the Src phosphorylation of emerin (Y74/Y95). Thus, emerin may protect nuclei from rupture when subjected to repeated pulling forces.

Here, we report that emerin activity is precisely calibrated in melanoma cells for fast amoeboid (leader bleb-based) migration. Using isolated nuclei, nuclei are found to be softest at intermediate levels of emerin activity. Surprisingly, Src activity is found to be lower in confined cells, as measured by a Fluorescence Resonance Energy Transfer (FRET) biosensor. Thus, nuclear deformability in melanoma cells may be enhanced by the down-regulation of Src.

## Results

Using human malignant melanoma, A375-M2 cells, we set out to determine how components of the nuclear lamina regulate fast amoeboid migration (Clark et al., 2000). To simulate the mechanical (compressive) forces in tissues, we subjected cells to a previously described assay (Jeremy et al., 2018). Briefly, cells are placed under a slab of PDMS, which is held at a defined height above cover glass by micron-sized beads. Both the PDMS and cover glass are coated with Bovine Serum Albumin (BSA; 1%), which provides friction for cell movement (Bergert et al., 2015). Using a range of cell types, it has been shown that confining cells down to 3 μm is the most effective at converting cells to fast amoeboid migration (Liu et al., 2015). Notably, the phenotypes observed *in vitro* are highly similar to what has been observed *in vivo* (Tozluoglu et al., 2013). Using this assay, we observe three distinct phenotypes, which we term leader mobile (LM), leader non-mobile (LNM), and no leader (NL) (Fig. 1A & Movies S1–3). The leader non-mobile (LNM) fraction represents cells that form leader blebs, which are stable over time, but do not move. No leader (NL) cells are also non-mobile. Interestingly, for all phenotypes the nucleus undergoes rapid shape changes (Fig. 1A; green channel). As a key component of the lamin nuclear skeleton, Lamin A levels have been shown to correlate with nuclear stiffness (Lammerding et al., 2006). Therefore, we wondered if increased levels of Lamin A would prevent nuclear membrane stretch and potential mechanosensing enzymes, such as cPLA2 (Lomakin et al., 2020; Venturini et al., 2020). In agreement with this concept, cells with increased Lamin A had nuclei that were rounder and were less frequently in leader blebs (Fig. 1B-C). The presence of the nucleus in leader blebs correlates with increased migration speed (Jeremy et al., 2018; Liu et al., 2015). Compared to EGFP alone, cells over-expressing mEmerald tagged Lamin A (Lamin A-mEmerald; Lamin A OE) were significantly slower (Fig. 1D). Moreover, when separated by phenotype, Lamin A OE cells were less likely to be mobile (Fig. 1E). However, the speed of mobile cells was unaffected by Lamin A OE (Fig. 1F). Therefore, the transition to the leader mobile (LM) phenotype was inhibited by Lamin A OE. Interestingly, Lamin A OE led to a ~50% reduction in leader bleb area (Fig. 1G). Lamin A was found to be ~1.5 fold over-expressed in transfected cells, as measured by Immunofluorescence (IF; Fig. 1H). In contrast, RNAi of Lamin A increased leader bleb area (Fig. S1A-H). These results are consistent with a model in which nuclear membrane stretch regulates actomyosin contractility through cPLA2. Thus, increasing levels of Lamin A correlate with increased nuclear stiffness and a reduction in motility (Fig. 1I). Using transmigration assays, we confirmed that the nuclear membrane tension senser, cPLA2, is required for confined migration in melanoma cells. More specifically, upon RNAi of cPLA2, transmigration through 8 μm pores is reduced by ~50%, whereas transmigration through 12 μm pores is unaffected (Fig. 1J).

**Figure 1.**
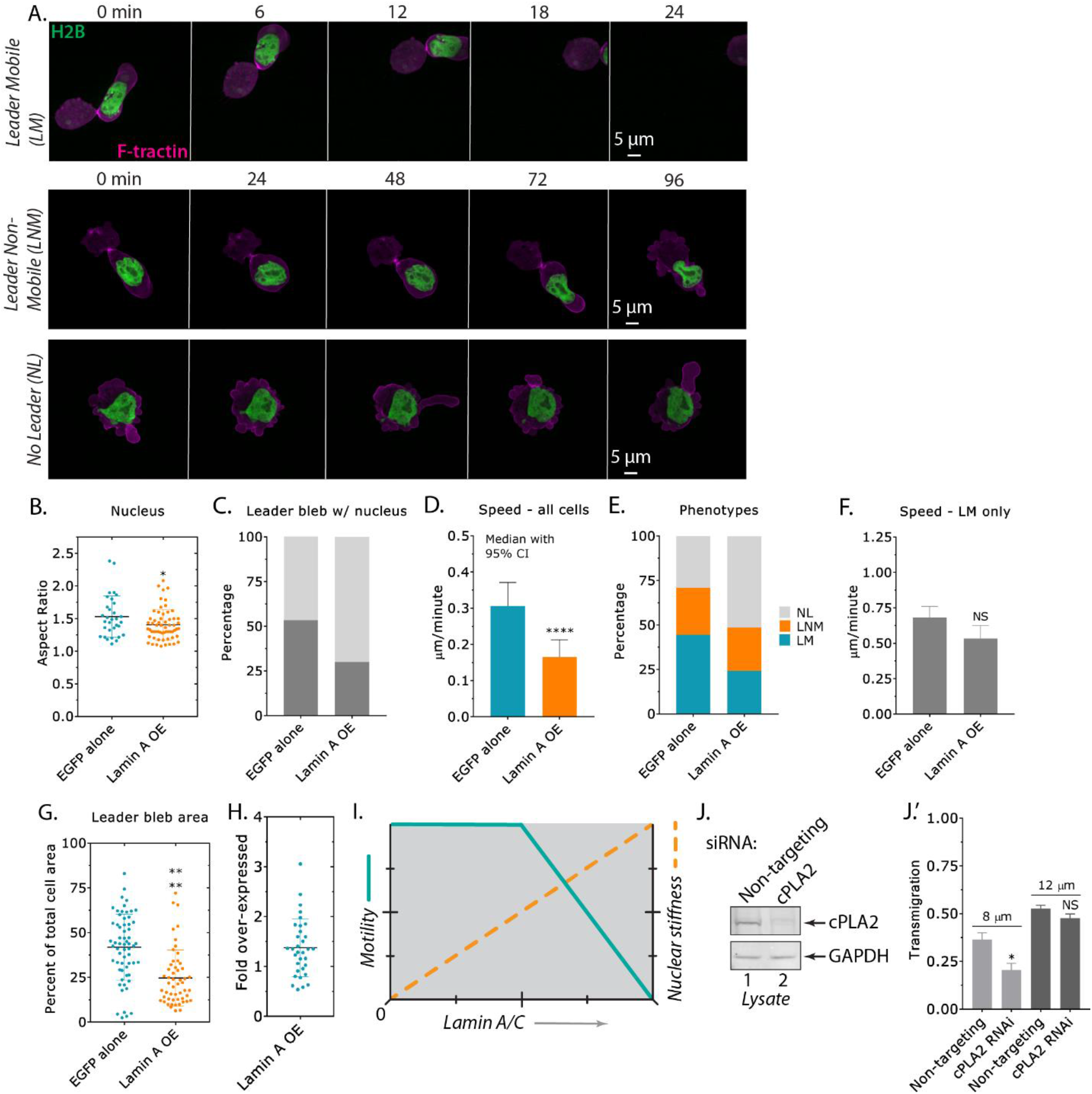
Fast amoeboid migration is inhibited by high levels of Lamin A. **A.** Time-lapse imaging of melanoma A375-M2 cells transiently transfected with H2B-mEmerald and F-tractin-FusionRed to mark nuclei and F-actin, respectively, which have been confined down to ~3 μm. **B.** Nuclear aspect ratio for cells with EGFP alone or Lamin A-mEmerald (mean +/- SD; two-tailed Student’s t-test). **C.** Percent cells with leader blebs containing the nucleus for EGFP alone or Lamin A-mEmerald. **D.** Instantaneous speeds for all cells with EGFP alone or Lamin A-mEmerald (median +/- 95% CI; two-tailed Student’s t-test). **E.** Percent NL, LNM, and LM for cells with EGFP alone or Lamin A-mEmerald. Statistical significance was determined using a Chi-squared test (χ^2^=1.16 x 10^-8^). **F.** Instantaneous speeds for leader mobile (LM) cells with EGFP alone or Lamin A-mEmerald (mean +/- SEM; two-tailed Student’s t-test). **G.** Leader bleb area (calculated as the percent of total cell area) for cells with EGFP alone or Lamin A-mEmerald (mean +/- SD; two-tailed Student t-test). **H.** Immunofluorescence (IF) was used to conduct a cell-by-cell analysis of Lamin A over-expression (OE; mean +/- SD). **I.** Motility and nuclear stiffness as a function of increasing Lamin A levels (conceptualized). **J.** Western blot confirming RNAi of cPLA2 in A375-M2 cells. **J.’** Transmigration of cPLA2 RNAi cells through 8 or 12 μm pores. All data are representative of at least three independent experiments. * - p ≤ 0.05, ** - p ≤ 0.01, *** - p ≤ 0.001, and **** - p ≤ 0.0001

The inner nuclear membrane (INM) protein, emerin, has been found to stiffen the nucleus in response to force (Guilluy et al., 2014). It has been proposed that emerin regulates nuclear stiffness through interactions with Lamin A (Berk et al., 2013). Initially, we wondered how cells would respond to the up- and down-regulation of emerin. Using EGFP tagged emerin (emerin-EGFP; emerin OE), we found emerin enriched at the nuclear envelope and ER of melanoma cells (Fig. 2A; green channel). Both when up- (emerin OE) and down-regulating (RNAi) emerin, nuclei are rounder and are found less often in leader blebs (Fig. 2B-C & S1A). Similarly, each condition led to significant decreases in cell speed and a smaller proportion of mobile cells (Fig. 2D-F). While RNAi of emerin led to a modest decrease in leader bleb area, emerin OE reduced leader bleb area by ~50% (Fig. 2G & Movie S4). Emerin was found to be ~2 fold over-expressed in transfected cells, as measured by IF (Fig. 2H). Collectively, these data suggest that fast amoeboid migration requires an intermediate (endogenous) level of emerin (Fig. 2I). Emerin RNAi led to a significant decrease in transmigration through 8 μm pores, whereas transmigration through 12 μm pores was unaffected (Fig. 2J). In emerin and cPLA2 RNAi cells, transmigration rates were reduced to a similar degree (Fig. 2K). Importantly, an additive effect on transmigration was not observed in emerin + cPLA2 RNAi cells, which suggests they are in the same pathway (Fig. 2K & 4A).

**Figure 2.**
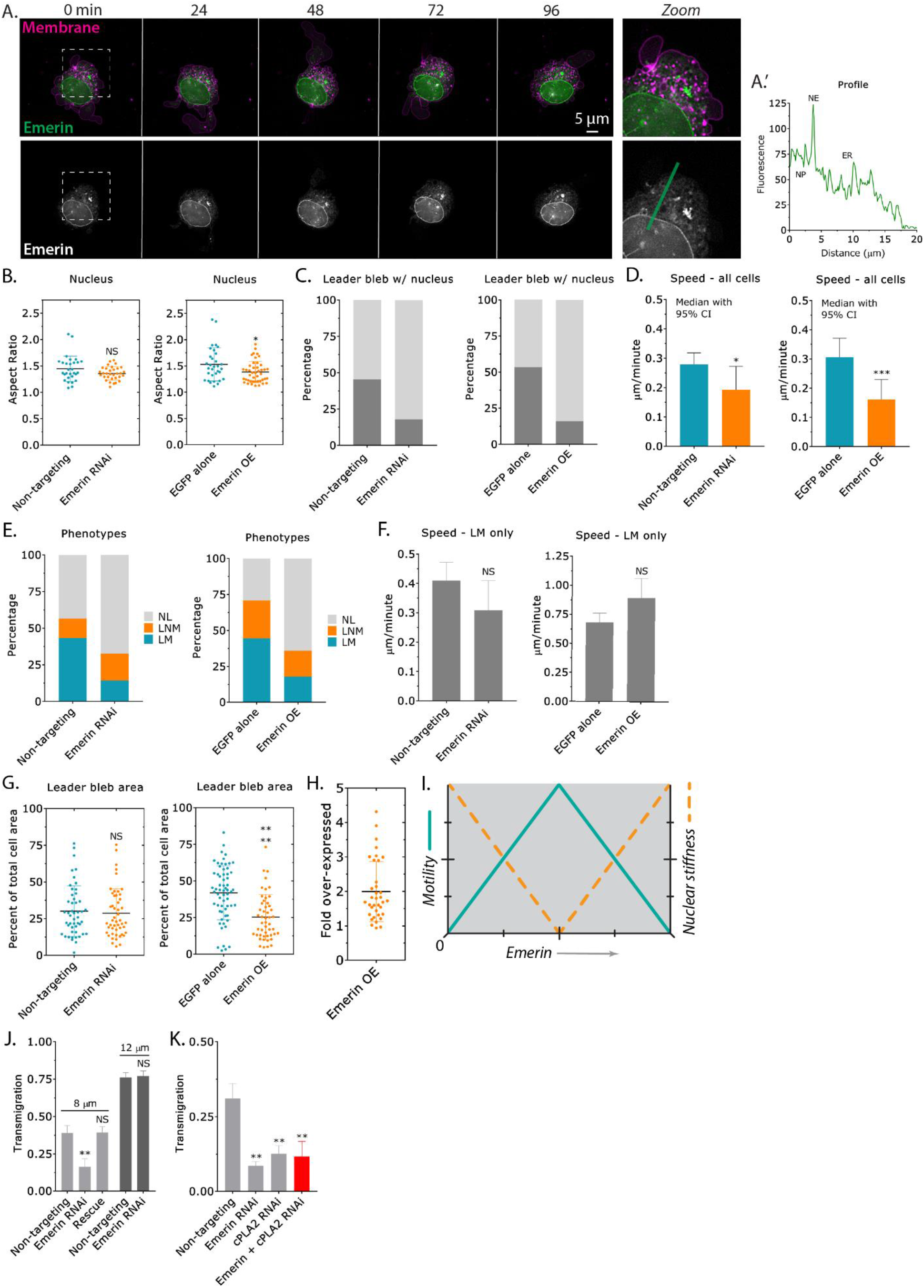
Fast amoeboid migration is inhibited by the up- or down-regulation of emerin. **A.** Time-lapse imaging of an A375-M2 cell transiently transfected with emerin-EGFP, which has been confined down to ~3 μm. Cells were stained with membrane dye. Zoom shows emerin predominantly at the nuclear envelope and ER. **A.’** Line-scan highlighting the enrichment of emerin at the nuclear envelope and ER. **B.** Nuclear aspect ratio of emerin RNAi (left) and over-expressing (OE; right) cells (mean +/- SD; two-tailed Student’s t-test). **C.** Percent cells with leader blebs containing the nucleus for emerin RNAi (left) and over-expressing (OE; right) cells. **D.** Instantaneous speeds for all emerin RNAi (left) and emerin over-expressing (OE; right) cells (median +/- 95% CI; two-tailed Student’s t-test). **E.** Percent NL, LNM, and LM for emerin RNAi (left) and over-expressing (OE; right) cells. Statistical significance was determined using a Chi-squared test (emerin RNAi; χ^2^=2.97 x 10^-7^, over-expression; χ^2^=7.69 x 10^-14^). **F.** Instantaneous speeds for leader mobile (LM) cells after emerin RNAi (left) and over-expression (OE; right) (mean +/- SEM; two-tailed Student’s t-test). **G.** Leader bleb area (calculated as the percent of total cell area) for emerin RNAi (left) and over-expressing (OE; right) cells (mean +/- SD; two-tailed Student’s t-test). **H.** IF was used to conduct a cell-by-cell analysis of emerin over-expression (OE; mean +/- SD). **I.** Motility and nuclear stiffness as a function of increasing emerin levels (conceptualized). **J.** Transmigration of emerin RNAi cells through 8 or 12 μm pores. An RNAi resistant version of emerin-EGFP was used in rescues. **K.** Transmigration of emerin, cPLA2, emerin + cPLA2 RNAi cells through 8 μm pores. All data are representative of at least three independent experiments. * - p ≤ 0.05, ** - p ≤ 0.01, *** - p ≤ 0.001, and **** - p ≤ 0.0001

In order to correlate our migration data with nuclear stiffness, we needed to measure nuclear stiffness. To accomplish this, we utilized a previously described approach, which involves sandwiching cells or isolated nuclei between two polyacrylamide (PA) gels of known stiffness (1 kPa). By taking the ratio of the height and width, we can measure “stiffness” (Liu et al., 2015). Using intact cells, neither emerin RNAi nor OE led to a significant change in stiffness (Fig. 3A-C). However, two emerin mutants, Phe240His-Frame Shift (FS; ΔINM) and Q133H, significantly increased cell stiffness (Fig. 3C). The Phe240His-FS mutation impairs the ability of emerin to localize to the inner nuclear membrane (INM), which has been previously described (Pfaff et al., 2016). The Q133H mutation is sufficient to cause Emery-Dreifuss Muscular Dystrophy (EMDM) and disrupts interactions between emerin and HDAC3, the pointed end of filamentous-actin (F-actin), and MAN1 (Berk et al., 2013). Through binding the pointed end, emerin is thought to stabilize F-actin near the INM (Holaska et al., 2004). In order to measure nuclear stiffness more directly, we disrupted cortical actin using Latrunculin-A (Lat-A; 500 nM). Using this approach, emerin RNAi led to a significant increase in stiffness (Fig. 3D). Although Lamin A OE increased stiffness, emerin OE led to a significant decrease in stiffness (Fig. 3E). Because of its interactions with F-actin, we wondered if F-actin may be important for emerin function. Therefore, we turned to using isolated nuclei. F-actin in the nucleus has been found to have numerous functions, including in DNA repair (Belin and Mullins, 2013). Using isolated nuclei, both emerin and Lamin A OE were found to increase stiffness (Fig. 3F). As demonstrated by emerin ΔINM, this effect is dependent on emerin being localized to the INM (Fig. 3F). Changes in nuclear aspect ratio, location, migration, and leader bleb area were similarly dependent on emerin being localized to the INM (Fig. S2A-I). Isolated nuclei with emerin Q133H were stiffer, but to a lesser degree when compared to emerin WT (Fig. 3F). Therefore, the Q133H mutation appears to not completely block nuclear stiffening. Likewise, nuclear aspect ratio, location, migration, and leader bleb area were intermediate between EGFP alone and emerin WT (Fig. S3A-H). The stiffness of isolated nuclei was unaffected by treatment with Lat-A, however, whether isolated nuclei are sensitive to actin monomer sequestering drugs is unknown (Fig. 3G). In contrast, de-condensation of chromatin with the histone deacetylase (HDAC) inhibitor, Trichostatin-A (3 μM), significantly decreased the stiffness of isolated nuclei (Fig. 3G). In parallel studies, using flow cytometry, the total level of F-actin in cells was found to be unaffected by emerin RNAi or OE (WT, ΔINM, Q133H, and Y74F/Y95F; Fig. S5A-B). Therefore, only when emerin is up-regulated, does nuclear stiffening require F-actin.

**Figure 3.**
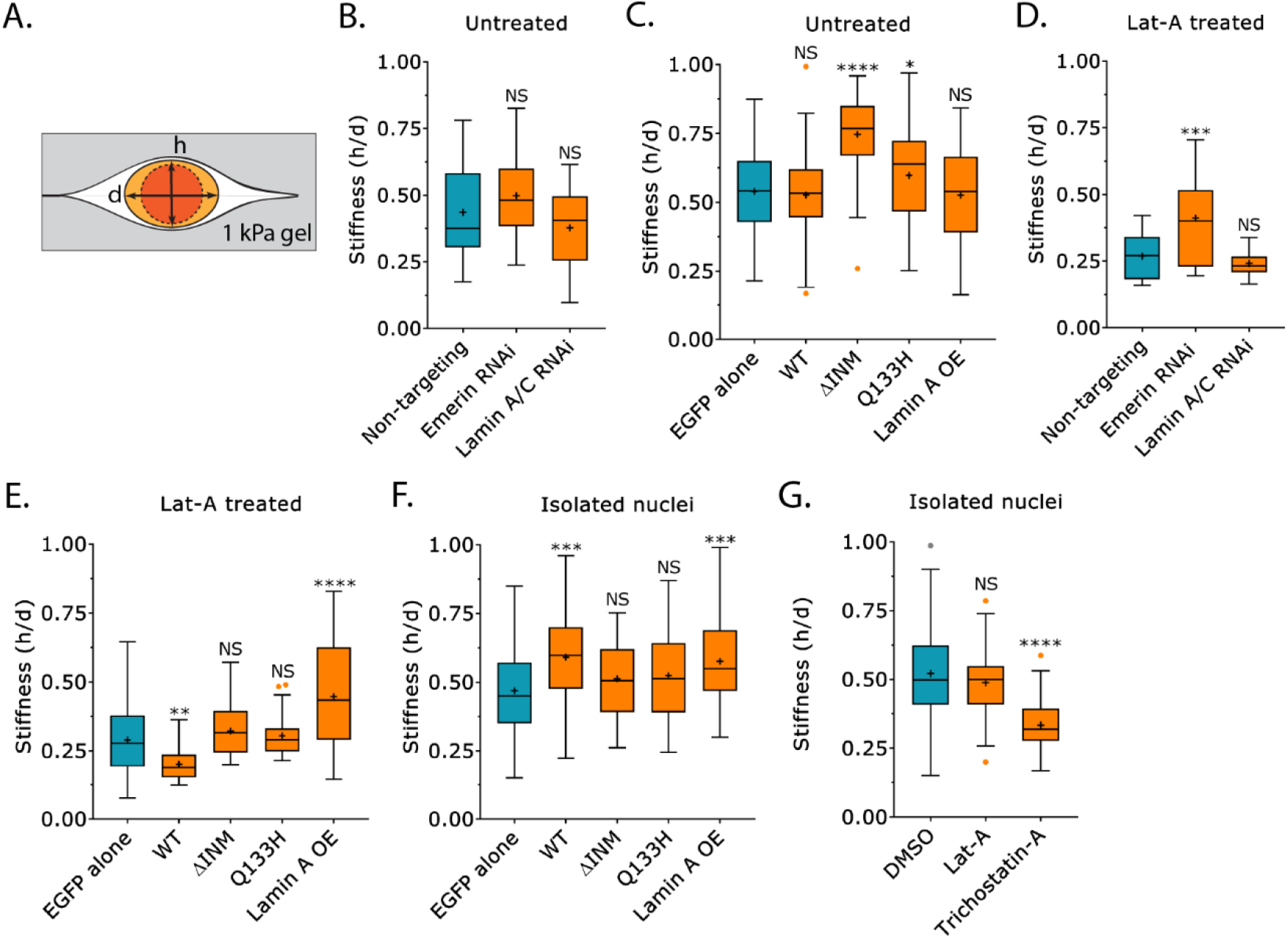
Nuclear stiffness is increased by the up- or down-regulation of emerin. **A.** Cartoon of the gel sandwich approach for measuring stiffness. **B.** Stiffness (*h/d*) for intact (untreated) cells after non-targeting (53 cells), emerin (49 cells), or Lamin A/C RNAi (46 cells) (“+” and line denote the mean and median, respectively; multiple-comparison test post-hoc). **C.** Stiffness (*h/d*) for intact (untreated) cells over-expressing (OE) EGFP alone (185 cells), emerin WT (70 cells), ΔINM (67 cells), Q133H (106 cells), and Lamin A (51 cells) (“+” and line denote the mean and median, respectively; multiple-comparison test post-hoc). **D.** Stiffness (*h/d*) for Latrunculin-A (Lat-A; 500 nM) treated cells after non-targeting (48 cells), emerin (54 cells), or Lamin A/C RNAi (48 cells) (“+” and line denote the mean and median, respectively; multiple-comparison test post-hoc). **E.** Stiffness (*h/d*) for Lat-A (500 nM) treated cells over-expressing (OE) EGFP alone (105 cells), emerin WT (47 cells), ΔINM (49 cells), Q133H (67 cells), and Lamin A (46 cells) (“+” and line denote the mean and median, respectively; multiple-comparison test post-hoc). **F.** Stiffness (*h/d*) for isolated nuclei (post cell fractionation) after over-expressing (OE) EGFP alone (105 nuclei), emerin WT (45 nuclei), ΔINM (45 nuclei), Q133H (41 nuclei), and Lamin A (57 nuclei) (“+” and line denote the mean and median, respectively; multiple-comparison test post-hoc). **G.** Stiffness (*h/d*) for isolated nuclei (post cell fractionation) after DMSO (166 nuclei), Lat-A (500 nM; 41 nuclei), and Trichostatin-A (3 μM; 90 nuclei) treatment (“+” and line denote the mean and median, respectively; multiple-comparison test post-hoc). All data are representative of at least three independent experiments. * - p ≤ 0.05, ** - p ≤ 0.01, *** - p ≤ 0.001, and **** - p ≤ 0.0001

It has been demonstrated that the Src mediated phosphorylation of emerin (Y74/Y95) is required for nuclei to stiffen in response to force (Guilluy et al., 2014). Therefore, we wondered if decreased fast amoeboid migration in emerin OE cells, which correlates with increased nuclear stiffness, is dependent on its phosphorylation (Fig. 4A). Cells with emerin (Y74F/Y95F) were found to have rounder nuclei, whereas the location of the nucleus was unaffected (Fig. 4B-D). Relative to emerin WT, cells with emerin (Y74F/Y95F) were significantly faster (Fig. 4E). Moreover, a larger fraction of cells with emerin (Y74F/Y95F) were mobile (Fig. 4F-G). Consistent with a role for nuclear stiffness in regulating fast amoeboid migration, while emerin WT increased the stiffness of isolated nuclei, the stiffness of isolated nuclei was unaffected by emerin (Y74F/Y95F; Fig. 4H-I). Moreover, treatment with the Src Family Kinase (SFK) inhibitor, Dasatinib (500 nM), significantly reduced the stiffness of isolated nuclei (Fig. 4J). In emerin (Y74F/Y95F) OE cells, leader bleb area was intermediate between EGFP alone and emerin WT (Fig. 4J & Movie S5). Emerin (Y74F/Y95F) was found to be ~2 fold over-expressed in transfected cells, as measured by IF (Fig. 2H). For cells over-expressing (OE) EGFP alone, emerin WT and Y74F/Y95F, F-actin and myosin were predominantly cortical in their localization (Fig. S4A-C). Accordingly, the inhibition of fast amoeboid (leader bleb-based) migration and increased nuclear stiffness by emerin OE requires its phosphorylation (Fig. 4L-M). Notably, the viability of cells after emerin RNAi and over-expression (OE) were similar (Fig. S5C-D).

**Figure 4.**
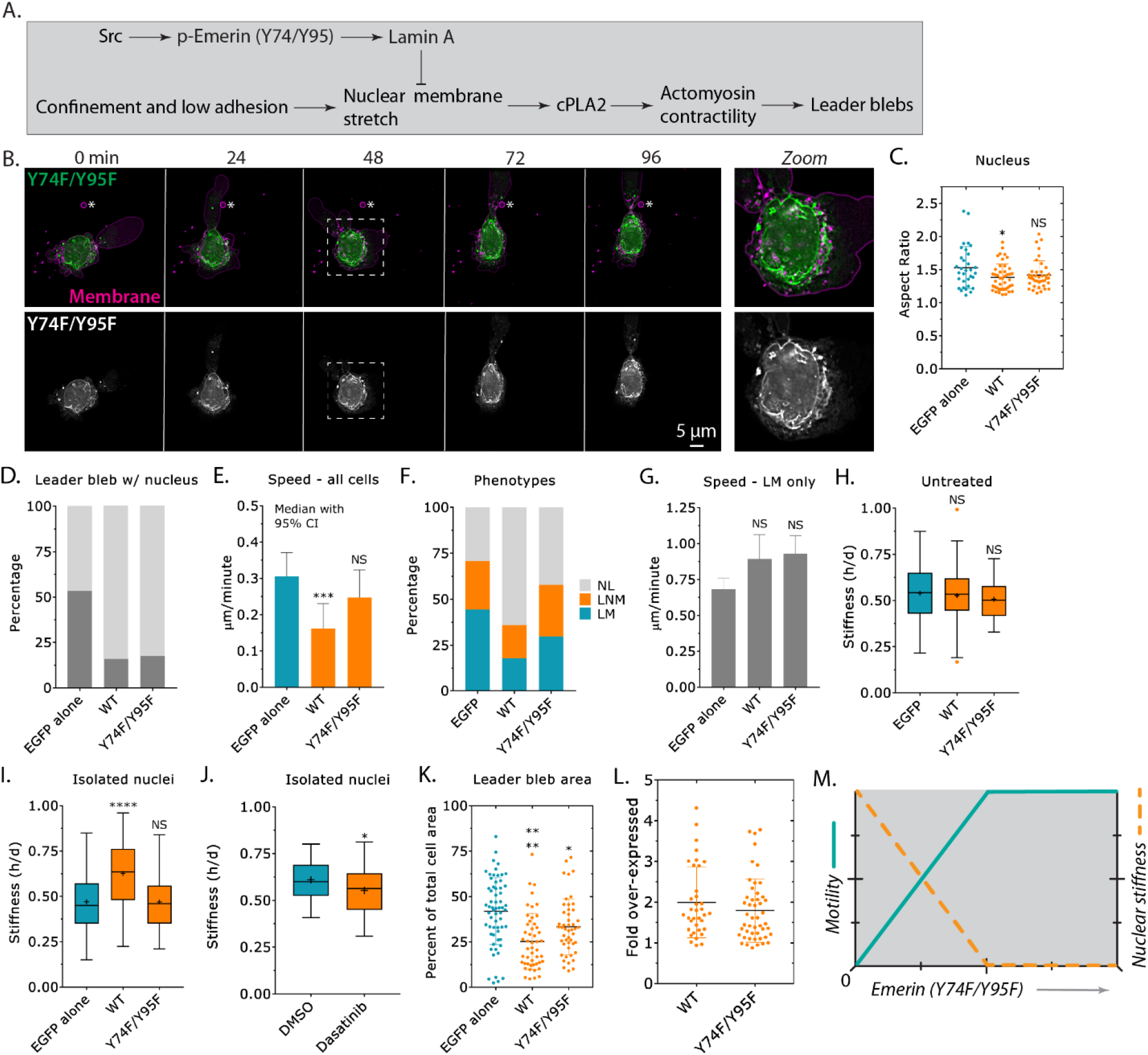
The Src mediated phosphorylation of emerin (Y74/Y95) inhibits fast amoeboid migration. **A.** Schematic of the pathway being investigated. **B.** Time-lapse imaging of a melanoma A375-M2 cell transiently transfected with emerin-EGFP (Y74F/Y95F), which has been confined down to ~3 μm. Cells were stained with membrane dye. Zoom shows emerin-EGFP (Y74F/Y95F) predominantly at the nuclear envelope and ER. **C.** Nuclear aspect ratio of cells over-expressing (OE) EGFP alone, emerin WT, and Y74F/Y95F (mean +/- SD; multiple-comparison test post-hoc). **D.** Percent cells with leader blebs containing the nucleus for EGFP alone, emerin WT, and Y74F/Y95F. **E.** Instantaneous speeds for all cells after over-expressing (OE) EGFP alone, emerin WT, and Y74F/Y95F (median +/- 95% CI; multiple-comparison test post-hoc). **F.** Percent NL, LNM, and LM for EGFP alone, emerin WT, and Y74F/Y95F. Statistical significance was determined using a Chi-squared test (EGFP alone vs. Y74F/Y95F; χ^2^=0.000104). **G.** Instantaneous speeds for leader mobile (LM) cells over-expressing (OE) EGFP alone, emerin WT, and Y74F/Y95F (mean +/- SEM; multiple-comparison test post-hoc). **H.** Stiffness (*h/d*) of intact (untreated) cells over-expressing (OE) EGFP alone (185 cells), emerin WT (70 cells), and Y74F/Y95F (57 cells) (“+” and line denote the mean and median, respectively; multiple-comparison test post-hoc). For comparison, data for EGFP alone and emerin WT were reproduced from figure 3C. **I.** Stiffness (*h/d*) of isolated nuclei (post cell fractionation) after over-expressing (OE) EGFP alone (81 nuclei), emerin WT (45 nuclei), and Y74F/Y95F (42 nuclei) (“+” and line denote the mean and median, respectively; multiple-comparison test post-hoc). For comparison, data for EGFP alone and emerin WT were reproduced from figure 3F. **J.** Stiffness (*h/d*) of isolated nuclei (post cell fractionation) after DMSO (52 nuclei) or Dasatinib (56 nuclei; 500 nM) treatment (“+” and line denote the mean and median, respectively; two-tailed Student’s t-test). **K.** Leader bleb area (calculated as the percent of total cell area) for cells over-expressing (OE) EGFP alone, emerin WT, and Y95F/Y74F (mean +/- SD; multiple-comparison test post-hoc). **L.** IF was used to conduct a cell-by-cell analysis of emerin WT and Y74F/Y95F over-expression (OE; mean +/- SD). **M.** Motility and nuclear stiffness as a function of increasing emerin (Y74F/Y95F) levels (conceptualized). All data are representative of at least three independent experiments. * - p ≤ 0.05, ** - p ≤ 0.01, *** - p ≤ 0.001, and **** - p ≤ 0.0001

Based on our data, we would predict that through the phosphorylation of emerin, high levels of Src activity would inhibit fast amoeboid migration (Fig. 4A). Therefore, we set out to determine the level of Src activity in confined cells. To accomplish this, we used a previously described Fluorescence Resonance Energy Transfer (FRET) based biosensor of Src activity (Ouyang et al., 2008). Briefly, this sensor has from its N- to C-terminus, a donor (ECFP), an SH2 domain, linker, a Src substrate peptide, acceptor (YPet), and a membrane anchor. Using freshly plated (spherical) cells, we confirmed that the emission ratio (acceptor/donor) of the sensor could be lowered by treatment with the SFK inhibitor, Dasatinib (0.5 μM; 10 min) (Fig. 5A). Relative to freshly plated (spherical) cells, the emission ratio (acceptor/donor) was significantly lower in cells confined down to ~3 μm (Fig. 5B). As an additional measure of Src activity, we calculated FRET efficiencies using the acceptor photobleaching method (Verveer et al., 2006). For freshly plated (spherical) cells, we calculated an average FRET efficiency of 17.29% (+/- 0.072; SD) (Fig. 5C-D). For cells confined down to 3 μm, we calculated an average FRET efficiency of 8.457% (+/- 0.032; SD), which is indicative of a decrease in Src activity in confined cells (Fig. 5C-D). Therefore, by down-regulating Src activity, melanoma cells appear to be primed for fast amoeboid (leader bleb-based) migration.

**Figure 5.**
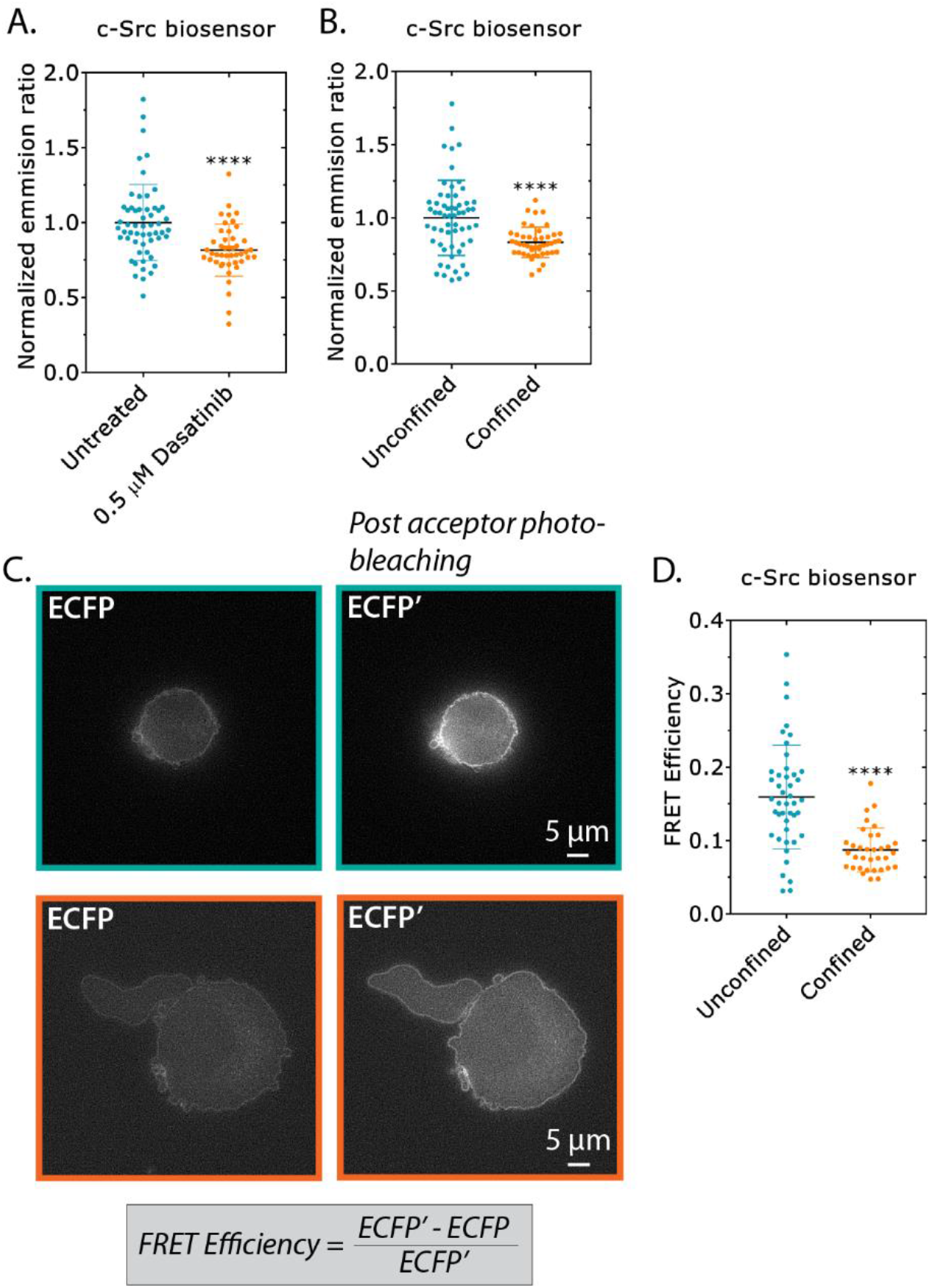
Confined cells have low levels of Src activity. **A.** Normalized emission ratio (acceptor/donor) for freshly plated (spherical) cells treated with the SFK inhibitor, Dasatinib (0.5 μM; 10 min) (mean +/- SD; two-tailed Student’s t-test). **B.** Normalized emission ratio (acceptor/donor) for freshly plated (spherical) cells and that have been confined down to 3 μm (mean +/- SD; two-tailed Student’s t-test). **C.** *Top*, ECFP (FRET donor) before and after acceptor photobleaching in a freshly plated (spherical) cell. *Bottom*, ECFP (FRET donor) before and after acceptor photobleaching in a cell that has been confined down to ~3 μm. The equation used for calculating FRET efficiencies is shown below. **D.** FRET efficiencies for freshly plated (spherical) cells and that have been confined down to ~3 μm (mean +/- SD; two-tailed Student’s t-test). All data are representative of at least three independent experiments. * - p ≤ 0.05, ** - p ≤ 0.01, *** - p ≤ 0.001, and **** - p ≤ 0.0001

## Discussion

As the largest organelle, the nucleus is subjected to an array of forces during cell migration. In confined environments, which may compress the nucleus, tension at the nuclear membrane with intracellular calcium was shown to activate cytosolic Phospholipase A2 (cPLA2) (Lomakin et al., 2020; Venturini et al., 2020). At the nuclear membrane, cPLA2 liberates arachidonic acid, elevating actomyosin contractility (Gong et al., 1992). Importantly, we could confirm using RNAi that cells require cPLA2 for transmigration through confining pores. Accordingly, the mechanical properties of the nucleus are likely to be a major regulator of fast amoeboid (leader bleb-based) migration.

In melanoma cells, we observed the nucleus undergoing rapid shape changes and moving into leader blebs during fast amoeboid migration. By up-regulating Lamin A, which is a key component of the lamin nuclear skeleton, we could suppress some of these shape changes and its movement into leader blebs (Ungricht and Kutay, 2017). Importantly, up-regulating Lamin A was found to inhibit the transition to fast amoeboid migration, which correlated with a decrease in leader bleb area. As Lamin A levels and nuclear stiffness correlate, these results suggest that the mechanical properties of the nucleus regulates fast amoeboid migration. It is noteworthy that down-regulating Lamin A was found to have a less pronounced effect. This may not be surprising, as many solid tumors already have low levels of Lamin A (Dubik and Mai, 2020).

Subsequently, we determined the role of the INM protein, emerin, in regulating fast amoeboid (leader bleb-based) migration. Although emerin loss of function mutations are well known in Emery-Dreifuss Muscular Dystrophy (EDMD), the role of emerin in regulating other cell biological processes is less well understood (Nagano et al., 1996). The discovery that emerin is key to nuclear stiffening in response to force, raises the possibility that emerin is critically important in diverse cell types. In motile cells, emerin may regulate the passage of cells through confined environments and protect the genome by preventing nuclear rupture (Denais et al., 2016; Raab et al., 2016). Accordingly, the perturbation of emerin function may prevent or abate cancer metastasis. In melanoma cells, we determined the role of emerin in regulating fast amoeboid migration. As in other cell types, emerin was predominantly localized to the nuclear envelope and ER in melanoma cells. Strikingly, up- or down-regulating emerin, led to rounder nuclei that were less often positioned in leader blebs. Cells in which emerin had been up- or down-regulated were also less often motile, correlating with a decrease in leader bleb area. Together, these results suggested that melanoma cells require intermediate or endogenous levels of emerin for fast amoeboid migration. In agreement with the concept that emerin regulates fast amoeboid migration through changes in nuclear membrane stretch, rates of transmigration through confining pores for RNAi of emerin, cPLA2, and emerin + cPLA2 were similar. However, as the nuclear lamina is known to play a role in chromatin organization, we cannot rule out the possibility that multiple genes are de-regulated as a result of changes in emerin activity.

Our results suggested that emerin may regulate fast amoeboid migration through changes in nuclear mechanics. Therefore, by measuring compressibility, we determined the stiffnesses of cells and isolated nuclei. Using intact cells, we were unable to detect any differences in stiffness after up- or down-regulating emerin. But after removing cortical actin with Lat-A, we could detect an increase in stiffness after down-regulating emerin. Surprisingly, in Lat-A treated cells, the stiffness of cells in which emerin was up-regulated was reduced. Therefore, we wondered if F-actin may be important for emerin function. Notably, emerin has been shown to interact with the pointed end of F-actin (Holaska et al., 2004). Indeed, using isolated nuclei from cells in which F-actin has not been perturbed, we could detect that emerin up-regulation increases stiffness. Moreover, this effect was dependent on emerin being localized to the INM. Interestingly, the stiffness of isolated nuclei with the Q133H mutant, which is found in EDMD patients, was intermediate between control (EGFP alone) and emerin WT (Berk et al., 2013). The Q133H mutation is known to disrupt interactions between emerin, HDAC3, F-actin, and MAN1 (Berk et al., 2013). This suggested to us that F-actin in the nucleus might regulate its stiffness, but we could not detect a significant decrease in stiffness when treating nuclei with Lat-A. However, actin monomer sequestering drugs may not be sufficient to disassemble F-actin in the nucleus. In contrast, chromatin de-condensation led to a significant reduction in the stiffness of isolated nuclei. In agreement with other labs, our data suggest that changes in chromatin structure are more likely to regulate nuclear stiffness (Nava et al., 2020).

In response to force, Src phosphorylates emerin (Y74/Y95), increasing nuclear stiffness (Guilluy et al., 2014). Therefore, we wondered if the inhibition of fast amoeboid migration by emerin OE is dependent on its phosphorylation. Indeed, we found that in cells with emerin (Y74F/Y95F), nuclei appeared more dynamic, as assessed by nuclear aspect ratio and localization. Moreover, cells were more often mobile, which correlated with softer isolated nuclei. Additionally, treating isolated nuclei with the SFK inhibitor, Dasatinib, led to a significant decrease in stiffness. Thus, high Src activity should inhibit fast amoeboid (leader bleb-based) migration. Accordingly, we determined the level of Src activity in confined cells. Strikingly, we found that relative to freshly plated (spherical) cells, confined cells had a lower level of Src activity. These data are consistent with previous work from our lab, which showed that fast amoeboid (leader bleb-based) migration is inhibited by high Src activity (Logue et al., 2018; Ullo and Logue, 2018). Apparently, when up-regulated, a significant fraction of emerin becomes phosphorylated even when Src activity is relatively low. Using isolated nuclei, it was reported that pulling anti-Nesprin-1 coated beads with magnetic tweezers activated Src; therefore, environments that require migrating cells to repeatedly pull on their nucleus (e.g., confining pores) may activate Src (Guilluy et al., 2014; McGregor et al., 2016). To the best of our knowledge, it is not known how pulling on Nesprin-1 activates Src near the nuclear envelope. Nesprin-1 connects F-actin in the cytoplasm to the nuclear lamina (Lombardi et al., 2011). As actin is predominantly cortical in fast amoeboid cells, the nucleus may not be subjected to strong pulling forces through Nesprin-1.

Collectively, our data demonstrate that the precise calibration of emerin activity is required for fast amoeboid (leader bleb-based) migration. Thus, the perturbation of emerin function may abate or prevent cancer metastasis.

## Supplemental Information

Supplemental information includes 5 figures and 5 movies and can be found with this article online.

## Methods

### Cell culture

A375-M2 cells (CRL-3223) were obtained from the American Type Culture Collection (ATCC; Manassas, VA). Cells were cultured in high-glucose DMEM (cat no. 10569010; Thermo Fisher, Carlsbad, CA) supplemented with 10% FBS (cat no. 12106C; Sigma Aldrich, St. Louis, MO), antibiotic-antimycotic (cat no. 15240112; Thermo Fisher), and 20 mM HEPES at pH 7.4 for up to 30 passages. Cells were plated at a density of 750,000 cells per well in a 6-well plate the day of transfection.

### Pharmacological treatments

Latrunculin-A (Lat-A; cat no. 3973), Trichostatin-A (cat no.1406), and Dasatinib (cat no. 6793) were purchased from Tocris Bioscience (Bristol, UK). All drugs were dissolved in DMSO (Sigma Aldrich). Cells were treated with Lat-A for 15 min, Trichostatin-A overnight, and Dasatinib for 30 min. Prior to cell stiffness measurements, polyacrylamide gels were incubated for 30 min in buffer S (20 mM HEPES at pH 7.8, 25 mM KCl, 5 mM MgCl_2_, 0.25 M sucrose, and 1 mM ATP) with DMSO or drug.

### Plasmids

F-tractin-FusionRed has been previously described (Logue et al., 2015). H2B-FusionRed was purchased from Evrogen (Russia). mEmerald-Lamin A-C-18 (Addgene plasmid no. 54138) and mEmerald-Nucleus-7 (Addgene plasmid no. 54206) were gifts from Michael Davidson (Florida State University). Emerin pEGFP-N2 (588; Addgene plasmid no. 61985) and Emerin pEGFP-C1 (637; Addgene plasmid no. 61993) were gifts from Eric Schirmer (University of Edinburgh). Kras-Src FRET biosensor (Addgene plasmid no. 78302) was a gift from Yingxiao Wang (University of Illinois). 1 μg of plasmid was used to transfect 750,000 cells in each well of a 6-well plate using Lipofectamine 2000 (5 μL; Thermo Fisher) in OptiMEM (400 μL; Thermo Fisher). After 30 min at room temperature, plasmid in Lipofectamine 2000/OptiMEM was then incubated with cells in complete media (2 mL) overnight.

### Mutagenesis

Emerin mutants were generated using the QuikChange II XL Site-Directed Mutagenesis Kit (Agilent Technologies; Santa Clara, CA) according to the manufacture’s protocol. The following forward primers were used for PCR:

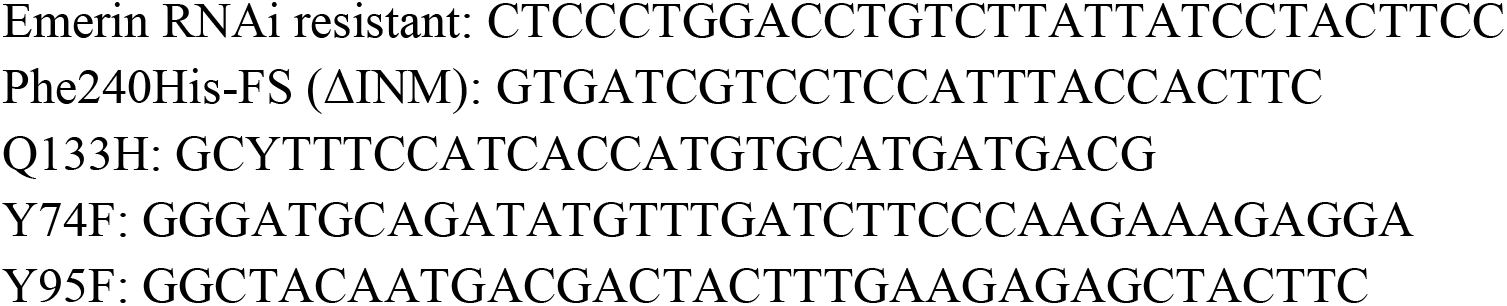

RNAi resistant emerin yields a single (silent; C->T) mutation centrally located within the LNA target sequence (GACCTGTCCTATTATCCTA). All clones were verified by sequencing using a commercially available resource (Genewiz, South Plainfield, NJ).

### LNAs

Non-targeting (cat no. 4390844), Lamin A/C (cat no. 4390824; s8221), and emerin (cat no. 4392420; s225840) Locked Nucleic Acids (LNAs) were purchased from Thermo Fisher. All LNA transfections were performed using RNAiMAX (5 μL; Thermo Fisher) and OptiMEM (400 μL; Thermo Fisher). Briefly, cells were trypsinized and seeded in 6-well plates at 750,000 cells per well in complete media. After cells adhered (~1 hr), LNAs in RNAiMAX/OptiMEM were added to cells in complete media (2 mL) at a final concentration of 50 nM. Cells were incubated with LNAs for as long as 5 days.

### Western blotting

Whole-cell lysates were prepared by scraping cells into ice cold RIPA buffer (50 mM HEPES pH 7.4, 150 mM NaCl, 5 mM EDTA, 0.1% SDS, 0.5% deoxycholate, and 1% Triton X-100) containing protease and phosphatase inhibitors (Roche, Switzerland). Before loading onto 4– 12% NuPAGE Bis-Tris gradient gels (Thermo Fisher), DNA was sheared by sonication and samples were boiled for 10 min in loading buffer. Following SDS-PAGE, proteins in gels were transferred to nitrocellulose membranes and subsequently immobilized by 0.1% glutaraldehyde (15 min) (Electron Microscopy Sciences; Hatfield, PA). After blocking in Tris-Buffered Saline containing 0.1% Tween 20 (TBS-T) and 1% BSA, primary antibodies against Lamin A/C (cat no. 2032; Cell Signaling Technology) or emerin (cat no. 30853; Cell Signaling Technology) were incubated with membranes overnight at 4 °C. Bands were then resolved with Horse Radish Peroxidase (HRP) conjugated secondary antibodies and a C-Digit imager (LI-COR Biosciences, Lincoln, NE). GAPDH (cat no. 97166; Cell Signaling Technology) was used to confirm equal loading.

### Immunofluorescence (IF)

After washing with HEPES-buffered saline (HBS), cells in 6-well glass-bottom plates (Cellvis) were fixed with 4% paraformaldehyde (Electron Microscopy Sciences) in HBS for 20 min at room temperature. Blocking, permeabilization, antibody incubation, and washing were done in HBS with 1% BSA, 1% fish gelatin, 0.1% Triton X-100, and 5 mM EDTA. A 1:250 dilution of Lamin A/C (cat no. 2032; Cell Signaling Technology) or emerin (cat no. 30853; Cell Signaling Technology) antibody was incubated with cells overnight at 4 °C. After extensive washing, a 1:400 dilution of Alexa Fluor 568–conjugated anti-rabbit secondary antibody (cat no. A-11036; Thermo Fisher) was then incubated with cells for 2 hr at room temperature. Cells were then incubated with a 1:1000 dilution of DAPI (cat no. D1306; Thermo Fisher). Cells were again extensively washed and then imaged in HBS. Cell-by-cell analyses of over-expression levels were conducted by comparing fluorescence intensities between transfected and untransfected cells. Transfected cells were identified using the fluorescent protein tag.

### Microscopy

High-resolution imaging was performed using a General Electric (Boston, MA) DeltaVision Elite imaging system mounted on an Olympus (Japan) IX71 stand with a computerized stage, Ultimate Focus, environment chamber (heat, CO_2_, and humidifier), ultrafast solid-state illumination with excitation/emission filter sets for DAPI, CFP, GFP, YFP, and Cy5, critical illumination, Olympus PlanApo N 60X/1.42 NA DIC (oil) objective, Photometrics (Tucson, AZ) CoolSNAP HQ2 camera, proprietary constrained iterative deconvolution, and vibration isolation table.

### Confinement

This protocol has been described in detail elsewhere (Jeremy et al., 2018). Briefly, PDMS (Dow Corning 184 SYLGARD) was purchased from Krayden (Westminster, CO). 2 mL was cured overnight at 37 °C in each well of a 6-well glass bottom plate (Cellvis, Mountain View, CA). Using a biopsy punch (cat no. 504535; World Precision Instruments, Sarasota, FL), an 8 mm hole was cut and 3 mL of serum free media containing 1% BSA was added to each well and incubated overnight at 37 °C. After removing the serum free media containing 1% BSA, 200 μL of complete media containing trypsinized cells (250,000 to 1 million) and 2 μL of beads (3.11 μm; Bangs Laboratories, Fishers, IN) were then pipetted into the round opening. The vacuum created by briefly lifting one side of the hole with a 1 mL pipette tip was used to move cells and beads underneath the PDMS. Finally, 3 mL of complete media was added to each well and cells were recovered for ~60 min before imaging. Here, complete media consisted of FluoroBrite (cat no. A1896701; ThermoFisher) supplemented with 20% FBS (cat no. 12106C; Sigma Aldrich), antibiotic-antimycotic (cat no. 15240112; Thermo Fisher), and 20 mM HEPES at pH 7.4.

### Leader blebs containing the cell nucleus

A thematic analysis of leader blebs and the nucleus was done for every cell by eye. If the nucleus moved into the leader bleb from the cell body and remained there for at least 3 consecutive frames, the cell was classified as having a ‘leader bleb containing the nucleus.’

### Classification of leader mobile (LM), leader non-mobile (LNM), and no leader (NL) cells

For classification, freshly confined cells were imaged every 8 min for 5 hr. Cells were classified by eye as either leader mobile (LM; cells that undergo fast directionally persistent leader bleb-based migration), leader non-mobile (LNM; cells with a leader bleb that persists for at least 5 consecutive frames but do not move), or no leader (NL; cells that undergo slow directionally non-persistent bleb-based migration).

### Instantaneous speed

Cells were tracked manually using the Fiji (https://fiji.sc/) plugin, MTrackJ, developed by Meijering and colleagues (Gorelik and Gautreau, 2014; Meijering et al., 2012). Instantaneous speeds from manual tracking were determined using the Excel (Microsoft; Redmond, WA) plugin, DiPer, developed by Gorelik and colleagues (Gorelik and Gautreau, 2014; Meijering et al., 2012). For minimizing positional error, cells were tracked every 8 min for 5 hr. Brightfield imaging was used to confirm that beads were not obstructing the path of a cell.

### Leader bleb area

For leader bleb area, freshly confined cells were imaged every 8 min for 5 hr and cells were traced from high-resolution images with the free-hand circle tool in Fiji (https://fiji.sc/). From every frame, the percent of total cell area for leader blebs was calculated in Excel (Microsoft) as the measured leader bleb area divided by the total cell area for the same frame. Measurements were then combined to generate an average for each cell.

### Cell stiffness assay

The previously described gel sandwich assay was used with minor modifications (Liu et al., 2015). 6-well glass bottom plates (Cellvis) and 18 mm coverslips were activated using 3-aminopropyltrimethoxysilane (Sigma Aldrich) for 5 min and then for 30 min with 0.5% glutaraldehyde (Electron Microscopy Sciences) in PBS. 1 kPa polyacrylamide gels were made using 2 μL of blue fluorescent beads (200 nm; ThermoFisher), 18.8 μL of 40% acrylamide solution (cat no. 161-0140; Bio-Rad, Hercules, CA), and 12.5 μL of bis-acrylamide (cat no. 161-0142; Bio-Rad) in 250 μL of PBS. Finally, 2.5 μL of Ammonium Persulfate (APS; 10% in water) and 0.5 μL of Tetramethylethylenediamine (TEMED) was added before spreading 9 μL drops onto treated glass under coverslips. After polymerizing for 40 min, the coverslip was lifted in PBS, extensively rinsed, and incubated overnight in PBS. Before each experiment, the gel attached to the coverslip was placed on a 14 mm diameter, 2 cm high PDMS column for applying a slight pressure to the coverslip with its own weight. Then, both gels were incubated for 30 min in media (cells) or buffer S (isolated nuclei) before plates were seeded. After the bottom gels in plates was placed on the microscope stage, the PDMS column with the top gel was placed on top of the cells seeded on the bottom gels, confining cells between the two gels (Fig. 3A). After 1 hr of adaptation, the height of cells was determined with beads by measuring the distance between gels, whereas the cell diameter was measured using a far-red plasma membrane dye (cat no. C10046; ThermoFisher). Stiffness was defined as the height (*h*) divided by the diameter (*d*). If drugs were used, gels were first incubated with drug in media for 30 min before an experiment.

### Nucleus isolation

The protocol for isolating nuclei has been previously described (Guilluy et al., 2014). 24 hr prior to nucleus isolation, 750,000 cells per well in a 6-well plate were transfected using Lipofectamine 2000 (5 μL; Thermo Fisher) as needed. Cells were lysed in 1 mL of hypotonic buffer (10 mM HEPES, 1 mM KCl, 1 mM MgCl_2_, 0.5 mM dithiothreitol, and protease inhibitors) for 10 min on ice. After cell fragments were detached using a scraper, samples were homogenized using 50 strokes of a tight-fitting Dounce homogenizer and then centrifuged at 700 x g for 10 min at 4 °C. Pellets were then washed in hypotonic buffer and centrifuged again. The nuclear pellet was then suspended in buffer S (20 mM HEPES at pH 7.8, 25 mM KCl, 5 mM MgCl_2_, 0.25 M sucrose, and 1 mM ATP) and then stained with a far-red fluorescent membrane dye (cat no. C10046; Thermo Fisher). Prior to the cell stiffness assay, the number and purity of nuclei as assessed by size was determined using an automated cell counter (TC20; Bio-Rad, Hercules, CA). If drug treated, drugs were present throughout the isolation process.

### Transmigration

Transmigration assays were performed using polycarbonate filters with 8 or 12 μm pores (Corning; Corning, NY). Prior to the assays, polycarbonate filters were coated with fibronectin (10 μg/mL; Millipore) then air dried for 1 hr. 100,000 cells in serum free media were seeded in the top chamber while the bottom chamber contained media with 20% FBS to attract cells. After 24 hr, cells from the bottom of the filter were trypsinized and counted using an automated cell counter (TC20; Bio-Rad). Transmigration was then calculated as the ratio of cells on the bottom of the filter over the total.

### Src biosensor

FRET efficiencies in unconfined vs. confined cells transfected with the Kras-Src FRET biosensor (Addgene plasmid no. 78302; gift from Yingxiao Wang) were calculated using the acceptor photobleaching method (Verveer et al., 2006).

### Flow cytometry

For measuring F-actin, trypsinized cells in FACS buffer (PBS with 1% BSA) were fixed using 4% paraformaldehyde (cat no. 15710; Electron Microscopy Sciences) for 20 min at room temperature. After washing, cells were stained with Alexa Fluor 647 conjugated phalloidin (cat no. A22287; Thermo Fisher) and DAPI (Sigma Aldrich) overnight at 4 °C. Data were acquired on a FACSCalibur (BD Biosciences; Franklin Lakes, NJ) flow cytometer. Median Fluorescence Intensities (MFIs) were determined using FlowJo (Ashland, OR) software. A detailed description of the flow gating strategy is shown in figure S5B.

### Statistics

All sample sizes were empirically determined based on saturation. As noted in each figure legend, statistical significance was determined by a two-tailed Student’s t-test or one-way ANOVA followed by a multiple-comparison test post-hoc. Normality was determined by a D’Agostino & Pearson test in Prism (GraphPad; San Diego, CA). * - p ≤ 0.05, ** - p ≤ 0.01, *** - p ≤ 0.001, and **** - p ≤ 0.0001

### Data availability

The data that support the findings of this study are available from the corresponding author, J.S.L., upon reasonable request.

## Acknowledgements

We thank members of the Logue Lab for insightful discussions. We would also like to thank the administrative staff within the Department of Regenerative and Cancer Cell Biology at the Albany Medical College. This work was supported by start-up funds from the Albany Medical College, a Young Investigator Award from the Melanoma Research Alliance (MRA; award no. 688232) (DOI: https://doi.org/10.48050/pc.gr.91570), and a Cancer Research Scholar Grant from the American Cancer Society (ACS; award no. RSG-20-019-01 - CCG) to J.S.L.

## Author Contributions

J.S.L. conceived and designed the study. S.B.L. performed all experiments except for transmigration assays (performed by A.P.C.), stiffness measurements on isolated nuclei treated with Dasatinib (performed by M.F.U.), and FRET assays (performed by J.S.L.). K.W.V. performed all flow cytometry and assisted with image analysis. A.K. provided technical assistance to S.B.L. J.S.L. wrote the manuscript with comments from S.B.L.

## SUPPLEMENTAL FIGURES

**Supplemental figure 1.**
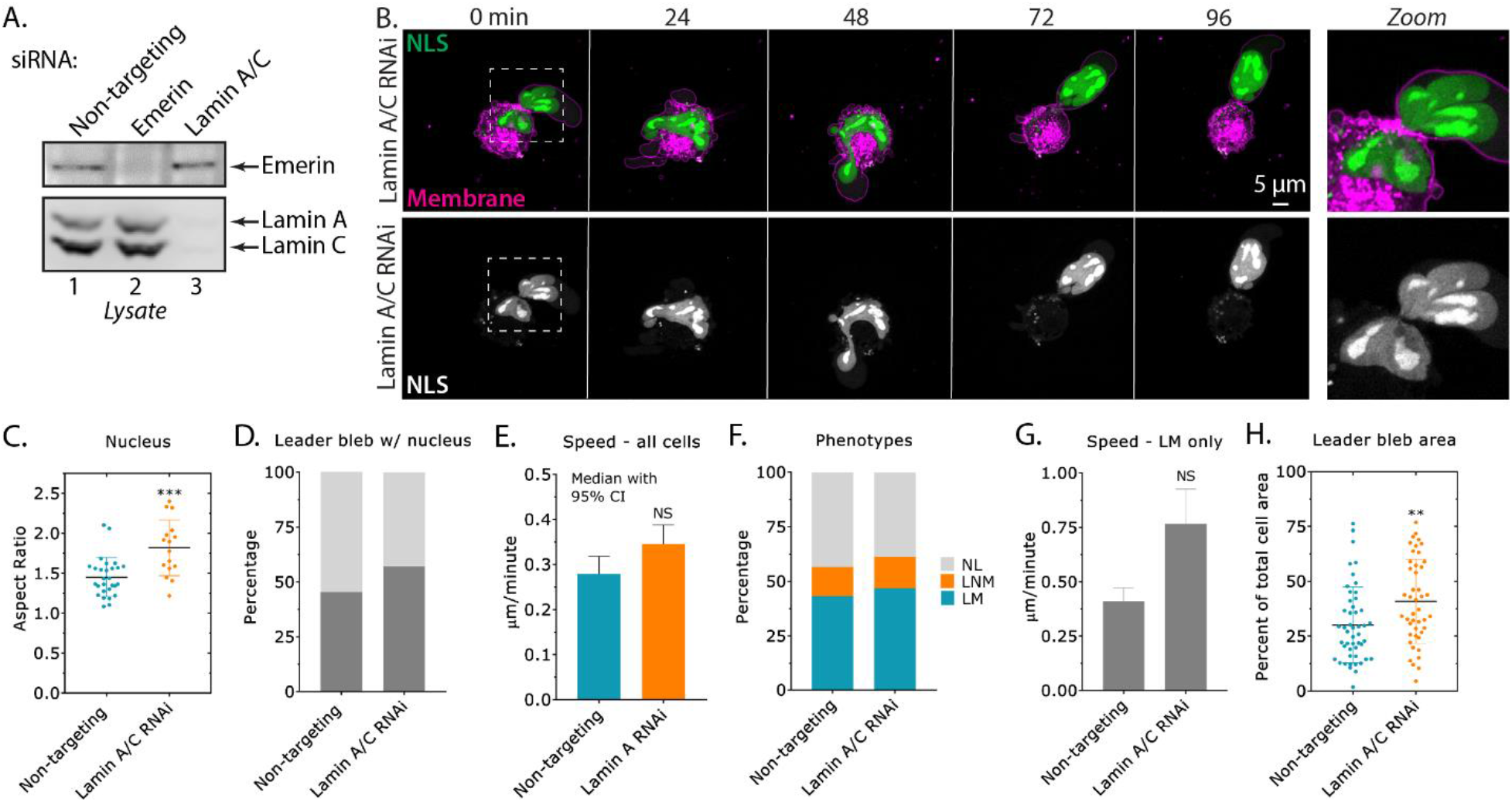
**A.** Western blot confirmation of emerin and Lamin A/C RNAi in A375-M2 cells. Cells were treated for 5 days with each Locked Nucleic Acid (LNA; 50 nM) to achieve a near complete removal of each protein. **B.** Time-lapse imaging of a A375-M2 cell after Lamin A/C RNAi transiently transfected with Nuclear Localization Sequence (NLS) tagged mEmerald (NLS-mEmerald), which has been confined down to ~3 μm. Cells were stained with membrane dye. Zoom highlights nuclear shape changes in Lamin A/C RNAi cells. **C.** Nuclear aspect ratio for cells after Lamin A/C RNAi (mean +/- SD; two-tailed Student’s t-test). **D.** Percent cells with leader blebs containing the nucleus in Lamin A/C RNAi cells. **E.** Instantaneous speeds for all cells after Lamin A/C RNAi (median +/- 95% CI; two-tailed Student’s t-test). **F.** Percent NL, LNM, and LM for Lamin A/C RNAi cells. Statistical significance was determined using a Chi-squared test (χ^2^=0.748). **G.** Instantaneous speeds for leader mobile (LM) cells after Lamin A/C RNAi (mean +/- SEM; two-tailed Student’s t-test). **H.** Leader bleb area (calculated as the percent of total cell area) for cells after Lamin A/C RNAi (mean +/- SD; two-tailed Student’s t-test). All data are representative of at least three independent experiments. * - p ≤ 0.05, ** - p ≤ 0.01, *** - p ≤ 0.001, and **** - p ≤ 0.0001

**Supplemental figure 2.**
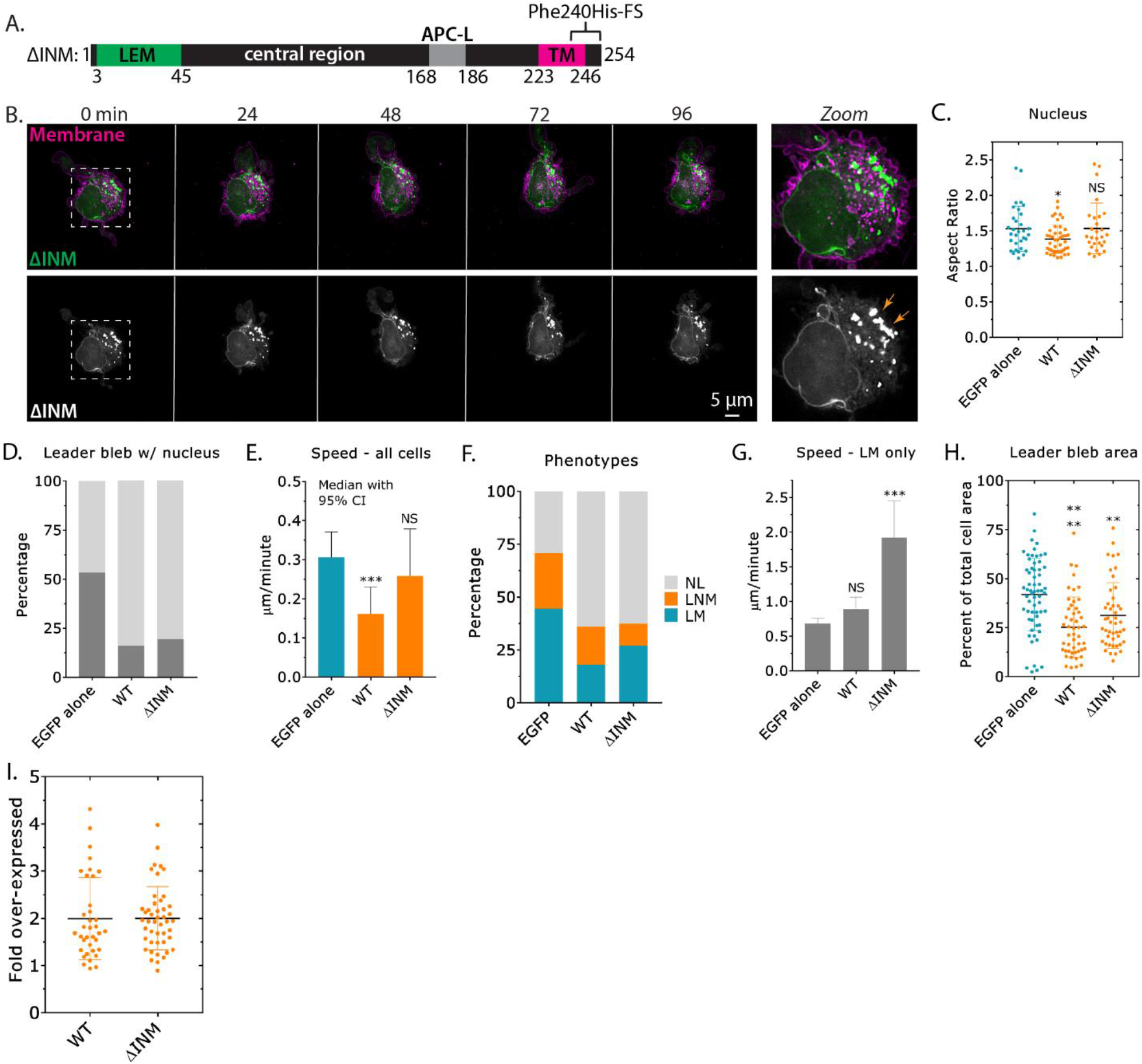
**A.** Cartoon of the previously described frame shift mutation, Phe240His-FS, near the transmembrane (TM) domain for retaining emerin within the ONM/ER (ΔINM). LEM (Lap2, emerin, MAN1) domain and APC-like (APC-L) domain. **B.** Time-lapse imaging of a A375-M2 cell transiently transfected with emerin-EGFP (ΔINM), which has been confined down to ~3 μm. Cells were stained with membrane dye. Zoom shows emerin (ΔINM) predominantly at the ONM/ER. **C.** Nuclear aspect ratio of cells over-expressing (OE) EGFP alone, emerin WT, and ΔINM (mean +/- SD; multiple-comparison test post-hoc). **D.** Percent cells with leader blebs containing the nucleus for EGFP alone, emerin WT, and ΔINM. **E.** Instantaneous speeds for all cells over-expressing (OE) EGFP alone, emerin WT, and ΔINM. **F.** Percent NL, LNM, and LM for EGFP alone, emerin WT, and ΔINM. Statistical significance was determined using a Chi-squared test (EGFP alone vs. ΔINM; χ^2^=2.42 x 10^-15^). **G.** Instantaneous speeds for leader mobile (LM) cells over-expressing (OE) EGFP alone, emerin WT, and ΔINM (mean +/- SEM; multiple-comparison test post-hoc). **H.** Leader bleb area (calculated as the percent of total cell area) for cells over-expressing (OE) EGFP alone, emerin WT, and ΔINM (mean +/- SD; multiple-comparison test post-hoc). **I.** Immunofluorescence (IF) was used to conduct a cell-by-cell analysis of emerin WT and ΔINM over-expression (OE; mean +/- SD). All data are representative of at least three independent experiments. * - p ≤ 0.05, ** - p ≤ 0.01, *** - p ≤ 0.001, and **** - p ≤ 0.0001

**Supplemental figure 3.**
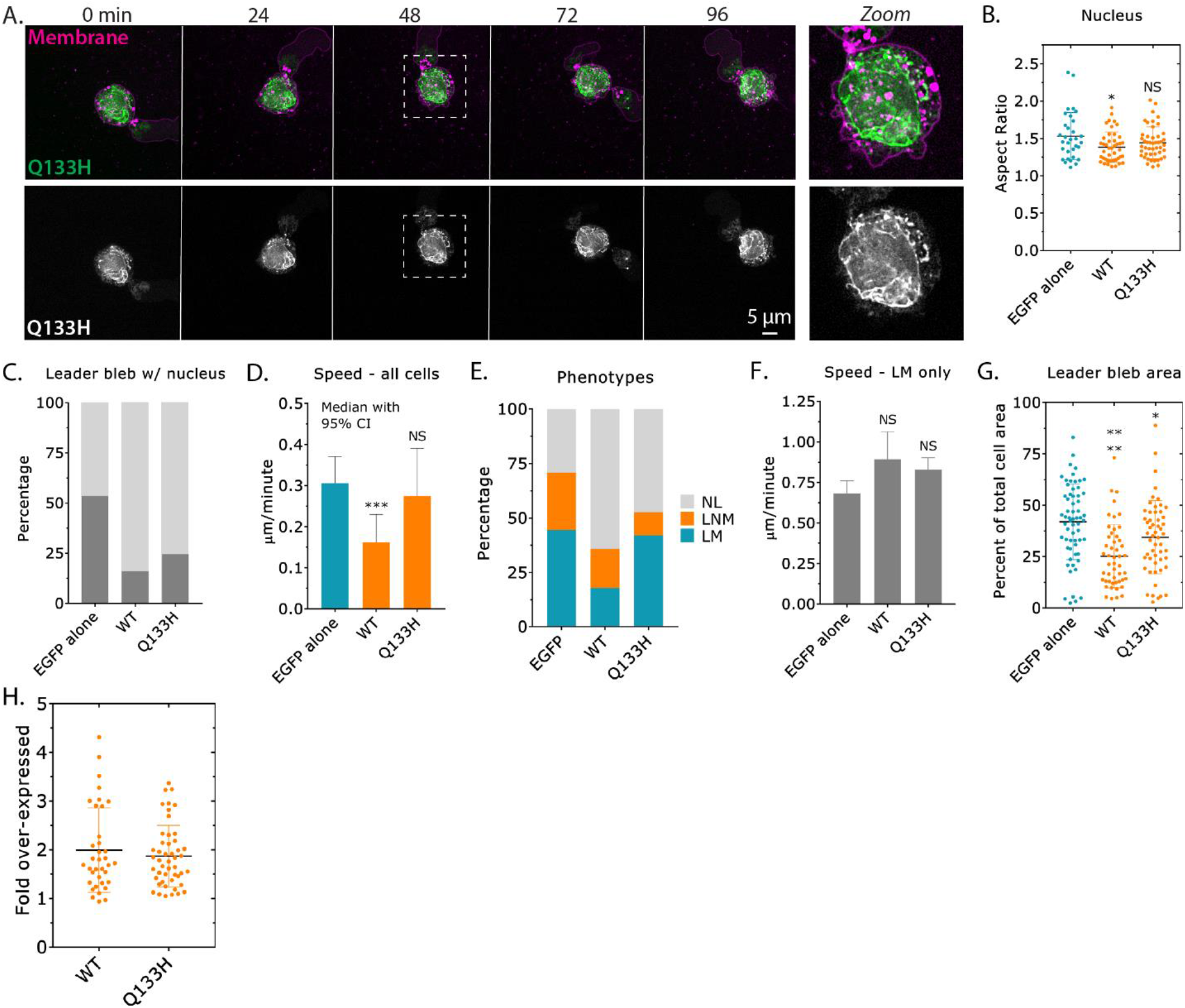
**A.** Time-lapse imaging of a A375-M2 cell transiently transfected with emerin-EGFP (Q133H), which has been confined down to ~3 μm. Cells were stained with membrane dye. Zoom shows emerin (Q133H) predominantly at the nuclear envelope and ER. **B.** Nuclear aspect ratio for cells over-expressing (OE) EGFP alone, emerin WT, and Q133H (mean +/- SD; multiple-comparison test post-hoc). **C.** Percent cells with leader blebs containing the nucleus for EGFP alone, emerin WT, and Q133H. **D.** Instantaneous speeds for all cells over-expressing (OE) EGFP alone, emerin WT, and Q133H (median +/- 95% CI; multiple-comparison test post-hoc). **E.** Percent NL, LNM, and LM for EGFP alone, emerin WT, and Q133H. Statistical significance was determined using a Chi-squared test (EGFP alone vs. Q133H; χ^2^=5.07 x 10^-6^). **F.** Instantaneous speeds for leader mobile (LM) cells over-expressing (OE) EGFP alone, emerin WT, and Q133H (mean +/- SEM; multiple-comparison test post-hoc). **G.** Leader bleb area (calculated as the percent of total cell area) for cells over-expressing (OE) EGFP alone, emerin WT, and Q133H (mean +/- SD; multiple-comparison test post-hoc). **H.** IF was used to conduct a cell-by-cell analysis of emerin WT and Q133H over-expression (OE; mean +/- SD). All data are representative of at least three independent experiments. * - p ≤ 0.05, ** - p ≤ 0.01, *** - p ≤ 0.001, and **** - p ≤ 0.0001

**Supplemental figure 4.**
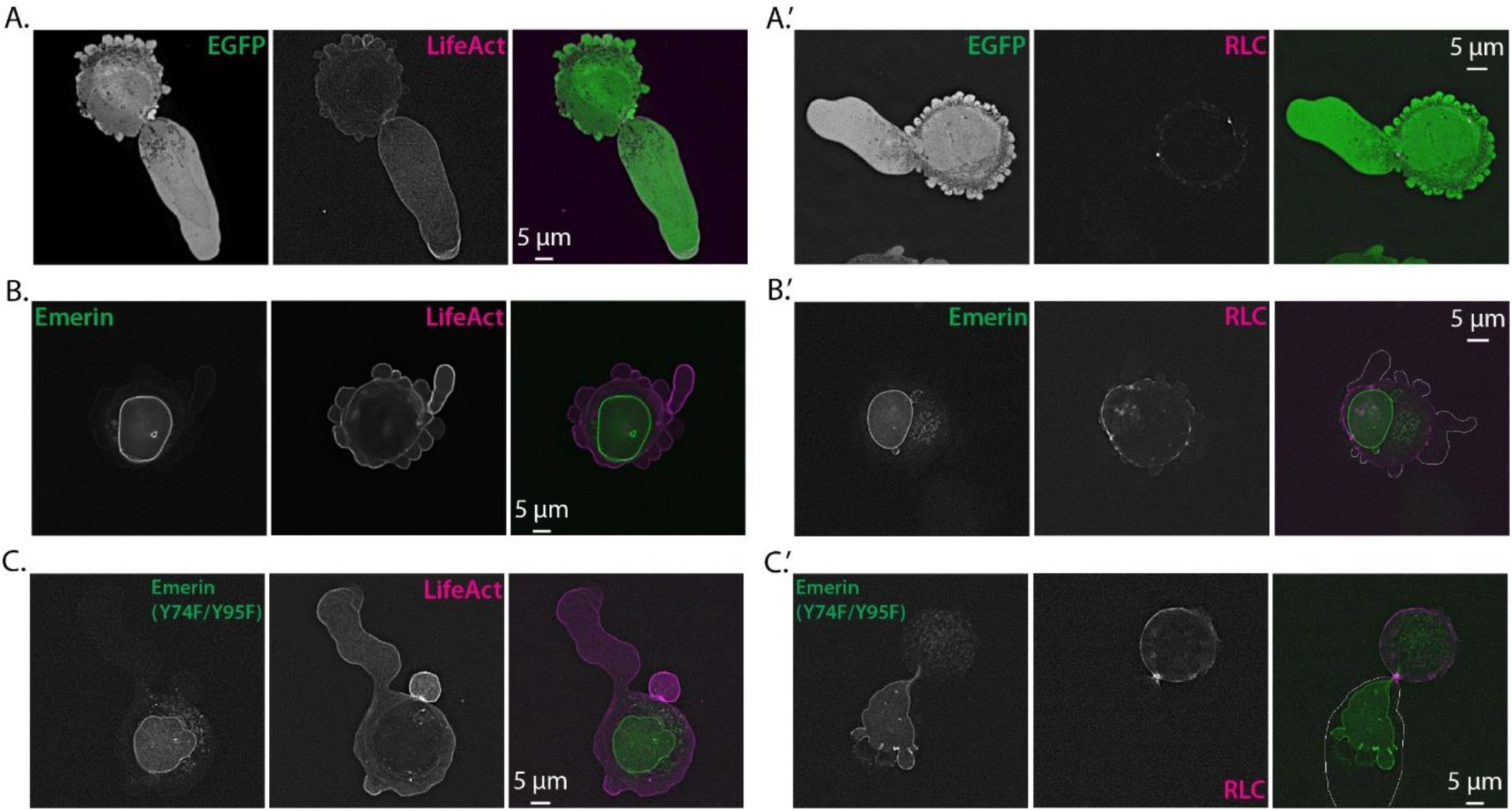
**A. - A.’** Representative images of an A375-M2 cell confined down to ~3 μm with (A) EGFP alone and mScarlet-LifeAct and (A’) EGFP alone and tdTomato-Regulatory Light Chain (RLC). **B. - B.’** Representative images of an A375-M2 cell confined down to ~3 μm with (B) emerin-EGFP and mScarlet-LifeAct and (B’) emerin-EGFP and tdTomato-RLC. The cell boundary was outlined in B’. **C. - C.’** Representative images of an A375-M2 cell confined down to ~3 μm with (C) emerin-EGFP (Y74F/Y95F) and mScarlet-LifeAct and (C’) emerin-EGFP (Y74F/Y95F) and tdTomato-RLC. The cell boundary was outlined in C’. All data are representative of at least three independent experiments.

**Supplemental figure 5.**
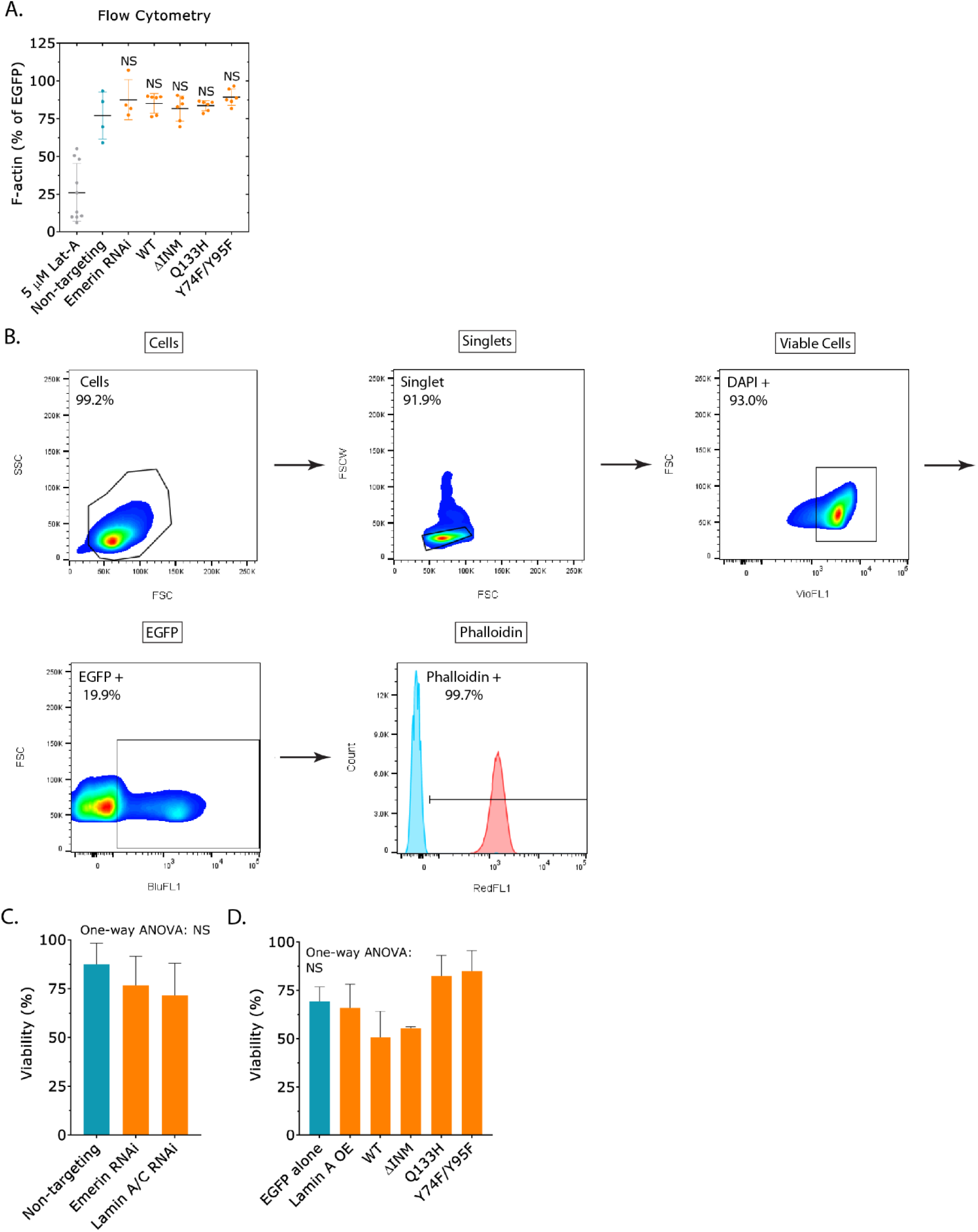
**A.** Flow cytometry analysis of total F-actin, as measured by far red fluorescent phalloidin binding in cells treated with Latrunculin-A (Lat-A; 5 μM), non-targeting, emerin RNAi, over-expressing (OE) emerin WT, ΔINM, Q133H, and Y74F/Y95F (relative to EGFP alone) (mean +/- SD; multiple-comparison test post-hoc). Each point corresponds to a Median Fluorescence Intensity (MFI). **B.** Flow gating strategy. Cells are isolated based on instrument calibration. Single cells were isolated by comparing forward scatter width and height. Viable cells were identified using DAPI staining while EGFP fluorescence was used to identify transfected cells. Cells that were positively stained with phalloidin (red curve) were isolated based on a gate created referencing unstained cells (blue curve) for analysis. **C.** Percent A375-M2 cells confined down to ∼3 μm alive after 5 hr of fluorescence imaging after non-targeting, emerin, and Lamin A/C RNAi (mean +/- SEM; One-way ANOVA). **D.** Percent A375-M2 cells confined down to ∼3 μm alive (viability) after 5 hr of fluorescence imaging over-expressing (OE) EGFP alone, Lamin A, emerin WT, ΔINM, Q133H, and Y74F/Y95F (mean +/- SEM; One-way ANOVA). All data are representative of at least three independent experiments. * - p ≤ 0.05, ** - p ≤ 0.01, *** - p ≤ 0.001, and **** - p ≤ 0.0001

## SUPPLEMENTAL MOVIES

**Supplemental movie 1.**
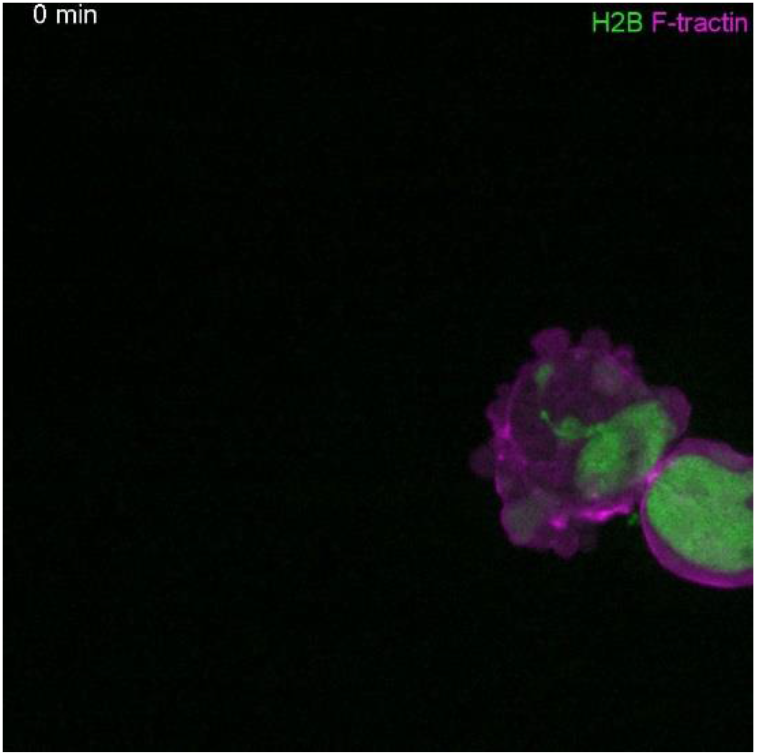
Time-lapse imaging of a leader mobile (LM) cell transiently transfected with H2B-mEmerald and F-tractin-FusionRed, which has been confined down to ~3 μm.

**Supplemental movie 2.**
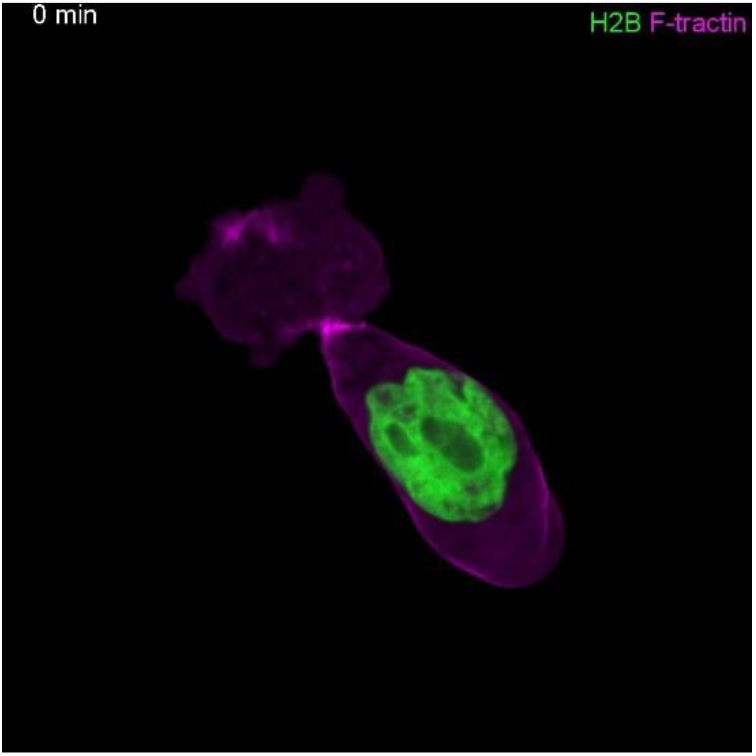
Time-lapse imaging of a leader non-mobile (LNM) cell transiently transfected with H2B-mEmerald and F-tractin-FusionRed, which has been confined down to ~3 μm.

**Supplemental movie 3.**
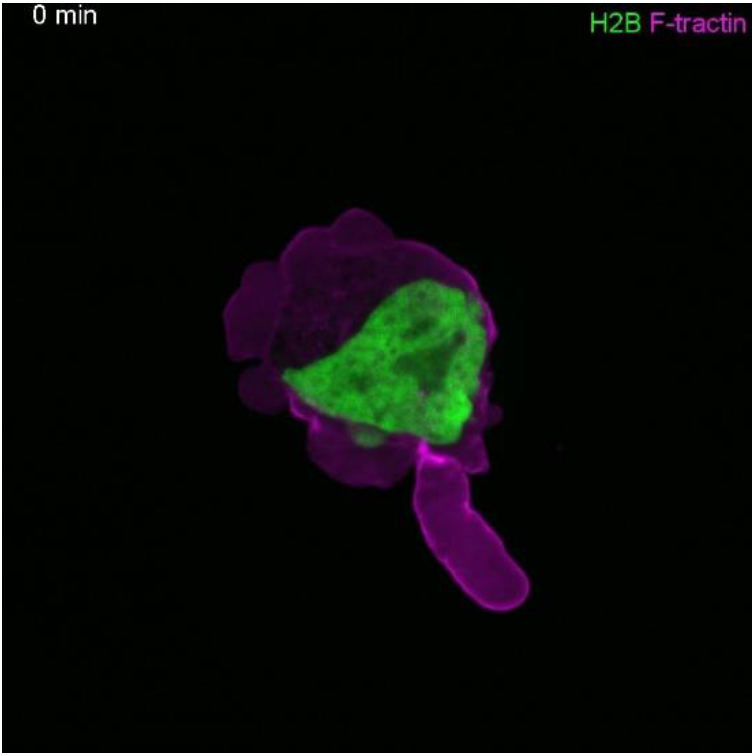
Time-lapse imaging of a no leader (NL) cell transiently transfected with H2B-mEmerald and F-tractin-FusionRed, which has been confined down to ~3 μm.

**Supplemental movie 4.**
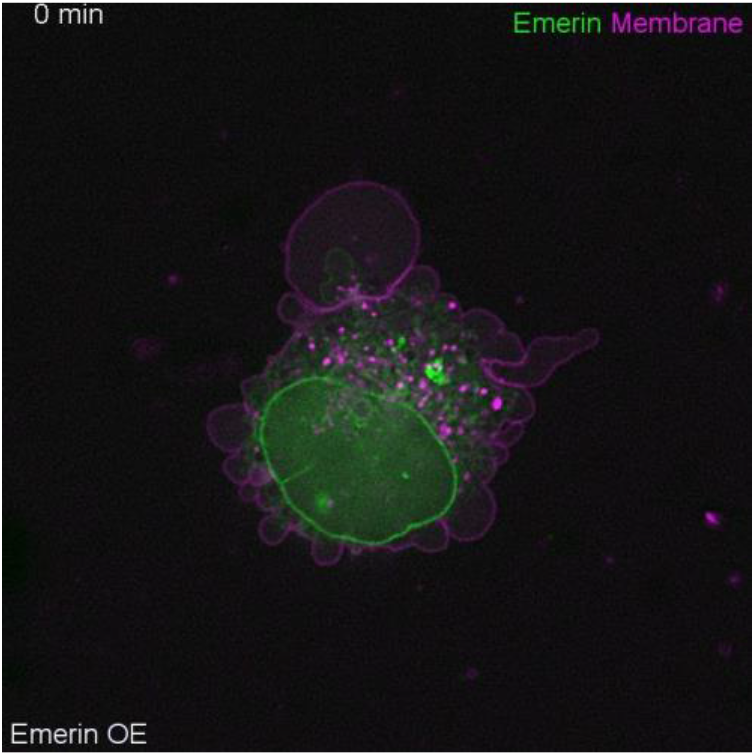
Time-lapse imaging of an A375-M2 cell over-expressing (OE) emerin-EGFP, which has been confined down to ~3 μm. Cells were stained with membrane dye.

**Supplemental movie 5.**
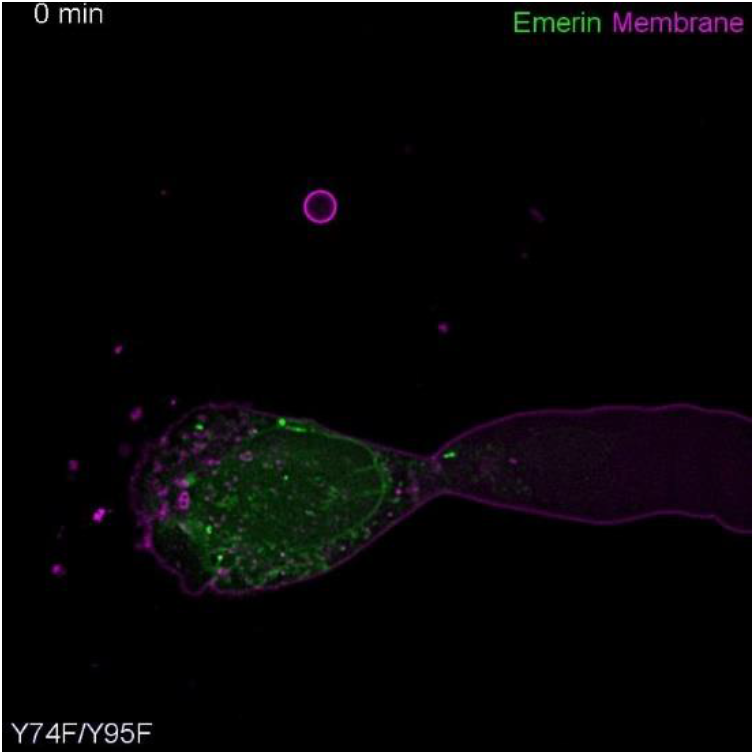
Time-lapse imaging of an A375-M2 cell over-expressing (OE) emerin-EGFP (Y74F/Y95F), which has been confined down to ~3 μm. Cells were stained with membrane dye.

